# Seed coating with phages for sustainable plant biocontrol of plant pathogens and influence of the seed coat mucilage

**DOI:** 10.1101/2024.01.12.575253

**Authors:** Sebastian H. Erdrich, Ulrich Schurr, Julia Frunzke, Borjana Arsova

## Abstract

Pathogens resistant to classical control strategies are on the rise and cause significant damage in crop yield production with seeds as one major transmission route. Bacteriophages are specialized viruses of bacteria and their interaction with seeds holds great potential as targeted and sustainable solution to this problem. In this study, we isolated and characterized two novel phages, Athelas and Alfirin, infecting *Pseudomonas syringae* and *Agrobacterium tumefaciens, respectively*, and included the recently published phage Pfeifenkraut infecting *Xanthomonas translucens*. The three phages were tested for their interaction with the seed coat mucilage. Phage binding on *Arabidopsis* seeds, which exude the mucilage as a polysaccharide-polymer-matrix, was assessed by comparison to seeds with removed mucilage. Two of the three phages were dependant on mucilage for seed binding, and podophage Athelas showed the highest dependency. Further podoviruses of the *Autographiviridae* obtained from the systematic *E. coli* (BASEL) phage collection were tested and showed a similar dependency on the mucilage for seed adhesion. Comparative analysis using a set of *Arabidopsis* seed coat mutants revealed the diffusible cellulose fraction as important component for phage binding. Long-term activity tests revealed a high stability of phages on seed surfaces and phage coating effectively increased the survival rate of plant seedling in the presence of the pathogen. Utilization of non-virulent host strains was further successfully applied to boost the presence of infectious phage particles on seed surfaces. Altogether, our study highlights the high potential of phage-based applications as sustainable biocontrol strategy on the seed level.

## 1. Introduction

Transmission of microbial diseases via seeds is a major concern in agriculture and can lead to considerable yield loss [^1–8^]. Some estimates predict that the usage of contaminated seeds can lead to yield reductions ranging from 15 to 90% [^9^]. This issue becomes especially critical in the face of a growing global population with an increasingly urgent demand for food, coupled with the looming threat of climate change that puts conventional agricultural methods’ productivity at risk. Bacteriophages as specialised viruses of bacteria could, in this context, offer a promising basis for the development of targeted and sustainable biocontrol strategies.

Phages have been discovered almost a century ago by d’Herelle and Twort and were used for first biocontrol trails shortly after that [^10^]. Nevertheless, with the discovery of broad range antibiotics, phages fell into oblivion due to their high specificity and lack of detailed knowledge which would have allowed to harness their full potential at that time. With the current rise of antibiotic or copper resistant bacteria classical methods to combat disease are getting less effective [^11–13^]. This has sparked a renaissance in phage research and moved them into the focus of researchers, policymakers and companies over the past decade [^14^]. In this context, biocontrol strategies centred around phages show significant promise, given the vast diversity of naturally occurring viruses [^15^].

In agriculture, different phage application methods have been explored recently, including spraying phage solutions on the phyllosphere [^16^], treatment of irrigation water in pot experiments [^17^], treating seed tubers [^18^] and coating leaves with formulations to protect the phages from radiation [^19^]. However, it has been reported by different studies that transmission via seeds appears as major route for plant pathogen transmission [^1–9^] and effective plant biocontrol via seed coating has been addressed by only a few studies in recent years. This includes for example the protection of melon plants from *Acidovorax citrulli* Rahimi-Midani by phage application [^20^] or the decontamination of seeds from *Xanthomonas campestris (Xcc)* [^21^]. Importantly, the establishment of effective phage coatings requires understanding of the binding mechanism, enrichment strategies and phage stability on seed surfaces, which has not been systematically addressed thus far.

One important seed product of many plant families is the seed coat mucilage, which is present in economically relevant plant families like *Lamiaceae* and *Solanaceae* as well as in the model plant *Arabidopsis thaliana* amongst many others [^22,23^]. The seed coat mucilage (SCM) is a layer of pectin, hemicelluloses, cellulose and proteins produced by the epidermal cells during seed development. It is released after imbibement of the mature seed with water and subsequently starts to swell and to cover the seed with a glycopolymer-matrix. Although the composition can differ amongst ecotypes of the same species, the major sugar-building-blocks are fucose, arabinose, rhamnose, galactose, glucose, mannose, xylose and galacturonic acid [^24^]. The mucilage was associated with anchorage to the soil and regulation of water content as well as an advantage in dispersal [^22,25^]. In this study, we assessed the influence of the seed coat mucilage on phage binding and stability.

Different phage morphotypes are characterized by their specific receptor binding proteins (RBP), encompassing tail fibers and tail spikes. These proteins play a crucial role in recognising chemical patterns on the surface of the host bacterium [^26,27^]. While certain receptors bind to the sugar moieties of polysaccharides, others target proteins in the cell envelope [^28^]. Consequently, phage adhesion to seeds may be facilitated by physical properties like the mesh-like polymer structure of the mucilage entrapping the phage particles, or through direct chemical interaction between the phage receptor binding proteins and specific sugar residues.

In this study, we systematically assessed phage binding and the relevance of the seed coat mucilage by focusing on the model plant *Arabidopsis thaliana* as well as representative bacterial plant pathogens. We describe two newly isolated phages infecting the prominent *Pseudomonas syringae* and *Agrobacterium tumefaciens* and included members from an *E*.*coli* phage collection in our tests [^29^]. While all phages tested showed binding to wild type *Arabidopsis* seeds, several phages showed significantly reduced binding to seeds with a removed mucilage layer. This included particularly phages of the *Autographiviridae* family, which were dependent on the presence of a mucilage. Testing several *Arabidopsis* seed mutants suggested a particular importance of the cellulose component of the mucilage. Further experiments confirmed a high stability of phages on seed surfaces without significant loss of infectivity. We are confident that this study will serve as an important step towards establishing future phage-based seed applications in agriculture.

## 2. Material and Methods

### 2.1 Bacterial strains and growth conditions

*Agrobacterium tumefaciens* (strain C58) and *Pseudomonas syringae* (DSM 50274) [^30^] were used as host strains for phage isolation in this study. *A. tumefaciens* cultures were grown on Lysogeny Broth (LB), whereas *P. syringae* (DSM 50274) and *Xanthomonas translucens* pv. translucens (DSM 18974) [^31^] cultures were grown on a nutrient agar. All cultures were inoculated from single colonies in the respective liquid media for overnight cultures. The cultivation of the bacterial strains was performed at 30 °C.

### 2.2 Phage isolation

The isolation of the phages was performed as previously described [^32^]. Briefly the virus particles within the environment samples were solubilized using phosphate-buffered saline (100 mM NaCl, 2.7 mM KCl, 10 mM Na_2_HPO_4_, 1.8 mM KH_2_PO_4_, 1 mM CaCl_2_, and 0.5 mM MgCl_2_; pH 7.5) and incubated for 3 h. Afterwards, the samples were centrifuged at 5000 *g* for 15 min to remove solid particles. The supernatants were filtered through 0.2 μm pore size membrane filters (Sarstedt; Filtropur S, PES). An aliquot of the filtered supernatant was plated on a 0.4% double agar overlay [^33^]. Plates were incubated at 30 °C overnight. Purification of the phage samples was carried out by re-streaking single plaques with an inoculation loop on a fresh double agar overlay for at least three times. The soil samples for *Agrobacterium* phage Alfirin were retrieved from the rhizosphere of a winter wheat plant at the IBG-2 crop garden (50.909277, 6.413403—Jülich, Germany) and phage Athelas was isolated from a wastewater sample donated by the Forschungszentrum Jülich wastewater plant (50.902547168169825, 6.404891888790708—Jülich, Germany). All phages will be available to the public via the German Collection of Microorganisms and Cell Cultures (DSMZ) after publication.

### 2.3 DNA isolation

Phage DNA was isolated by treating 2 ml of Phage solution with 1 U/μL DNAse (Invitrogen, Carlsbad, CA, USA) to remove free DNA. The further steps of DNA isolation were performed with the Norgen Biotek Phage DNA Isolation Kit (Norgen Biotek, Thorold, Canada) according to the manufacturers protocol.

### 2.4 DNA sequencing and genome assembly

Assembly the DNA library was performed using the NEBNext Ultra II DNA Library Prep Kit for Illumina, according to the manufacturer’s instructions, and shotgun-sequenced using the Illumina MiSeq platform with a read length of 2 × 150 bp (Illumina, San Diego, CA, USA). For each phage, a subset of 100,000 reads were sampled, and a de novo assembly was performed using CLC genomics workbench 20.0.4 (QIAGEN, Hilden, Germany). Finally, the obtained contigs were manually curated and checked for gene coverage.

### 2.5. Gene prediction and functional annotation

The phage open reading frames (ORFs) were predicted with Pharokka v 1.3.2 [^34^] in terminase reorientation mode using PHANOTATE [^35^], tRNAs were predicted with tRNAscan-SE 2.0 [^36^], tmRNAs were predicted with Aragorn [^37^] and CRISPRs were checked with CRT [^38^]. Functional annotation was generated by matching each CDS to the PHROGs [^39^], VFDB [^40^] and CARD [^41^] databases using MMseqs2 [^42^] and PyHMMER [^43^]. Contigs were matched to their closest hit in the INPHARED database [^44^] using mash [^45^]. Plots were created with the pyCirclizen package. Additionally, all identified sequences were later curated usually manually using online NCBI Blast against the non-redundant (NR) database [45]. Conserved protein domains were further predicted using the batch function of NCBI Conserved Domain Database (CDD) [46] with the e-value cut-off of 0.01.

### 2.6. Electron microscopy of phage virions

For electron microscopy of single phage particles, 3.5 μL purified phage suspension was fixated on a glow discharged (15 mA, 30 s) carbon coated copper grid (CF300-CU, carbon film 300 mesh copper) and stained with 2% (w/v) uranyl acetate. After air drying, the sample was analyzed with a TEM Talos L120C (Thermo Scientific, Dreieich, Germany) at an acceleration of 120 kV.

### 2.7. Sterilization of *Arabidopsis thaliana seeds*

The seed coat was sterilised by vortexing for 5 min in 50% ethanol (EtOH) containing 0.5% Triton x-100. Afterwards, the 50% EtOH was removed and replaced by 96% EtOH, the samples were inverted once. Afterwards, all EtOH was removed immediately. The seeds were transferred within a small volume of 96% EtOH onto sterile filter paper using a pipette. Finally, they were air dried.

### 2.8 Phage binding to wild type seeds of *Arabidopsis thaliana*

Approximately 1000 *Arabidopsis thaliana* Col-0 seeds were incubated in a sterile Eppendorf tube with 1 mL bacteriophage solution of a concentration of 10^8^ Pfu/mL or higher for 30 minutes. Followed by two subsequent washing steps in ddH_2_O to remove non-bound phage particles. Afterwards the seeds were air-dried on filter paper beneath a clean bench until the liquid was evaporated for 30-45 min.

### 2.9 Influence of seed coat mutants on phage binding

The following seed coat mutants were required from Nottingham Arabidopsis Stock Centre (NASC): ttg1-21 (GK-580A05); csla2-3 (SALK_149092); rhm2 (SALK_076300); muci70-1 (SALK_129524). The cesa5 and sbt 1.7 mutant were present at the IBG-2. All seeds were germinated on sterile ½ MS plates and subsequently propagated on soil using a 16h day/ 8h night regime.

Afterwards the harvested seeds were checked using ruthenium red staining [^46^] for their typical morphological appearance under the microscope. The seeds were subsequently used for phage binging assays as described above.

For mechanical removal of the mucilage layer (Col-0 mucilage extracted), approx. 1000 EtOH sterilised wild-type seeds were incubated for 5 minutes in 1 mL ddH_2_O, followed by 2 rounds of 15 min 30Hz/s shaking in a ball-mill (Retsch MM200, Retsch, Germany) without beads [^46^]. The mucilage containing supernatant was removed and the seeds washed in in two subsequent steps with ddH_2_O. Afterwards, the seeds were placed on sterile filter papers and air-dried for 30 min under the clean bench. Complete removal of the mucilage was verified by ruthenium red staining and microscopy before further use.

### 2.10 Phylogeny of PRB Proteins and in silico protein folding

Ancestral states were inferred using the Maximum Likelihood method [^47^] and JTT matrix-based model [^48^]. The tree (Fig 5.) shows a set of possible amino acids (states) at each ancestral node based on their inferred likelihood at site 1. Initial tree(s) for the heuristic search were obtained automatically by applying Neighbor-Join and BioNJ algorithms to a matrix of pairwise distances estimated using the JTT model, and then selecting the topology with superior log likelihood value. The rates among sites were treated as being uniform among sites (Uniform rates option). This analysis involved 24 amino acid sequences. There were a total of 1386 positions in the final dataset. 1000 Bootstrap trees were generated for the final tree. Evolutionary analyses were conducted in MEGA11 [^49^].

The 3D protein structures of phage receptor binding proteins were predicted using the ColabFold v1.5.3 webserver with AlphaFold2 using MMseqs2 with the default settings[^50^].

### 2.11 Stability of phages on plant seeds

To evaluate the stability of phages on seeds the *Agrobacterium phage Alfirin*, the *Pseudomonas phage Athelas* and the *Xanthomonas phage Pfeifenkraut*[^32^] were bound to *Arabidopsis thaliana Col-0* seeds as described above and stored within microcentrifuge tubes at 4°C for up to 28 days. At defined timepoints a portion of the seeds taken from the tubes and incubated at 28°C for 24 hours on a double agar overlay containing the respective host-bacterium. Finally, the total amount of seeds as well as the portion showing plaques around the seeds were counted.

### 2.12 Survival of seedlings in presence of the pathogen

The survival in presence of the pathogen was assessed by infecting sterilised Col-0 seeds artificially by imbibing them in a bacterial solution of *Agrobacterium tumefaciens C58*at an OD of 0.4 for 30 min and subsequent air-drying on sterile filter paper. The same procedure was used for the phage as described above. All seeds were sown on ½ MS Agar plates and placed into the climate chamber with 12/12h day/night regime at 22°C at day and 19°C at night. Scans of the plates were taken at 14 days after sowing to evaluate the survival rate. All germinated seedlings surpassing the 2-cotyledon stage without signs of necrosis were counted as alive.

### 2.13 Booster with a nonvirulent host

The “Booster” experiment, aimed to investigate incubation with non-virulent strains as a strategy for the targeted enrichment of phages at the plant seeds. Therefore, the experiment was performed with non-virulent agrobacterium[^51^] and phage co-coated on Col-0 seeds with 1*10^9^ cfu/mL. Phage Alfirin was used for this experiment at a MOI of 1. The coating was performed as described above for the phages. The seeds were placed ½ MS Agar the plates were incubated for 14 days within the climate chamber. Subsequently the seedlings were transferred to a LB medium based double agar overlay, containing the wildtype *Agrobacterium tumefaciens*, at an OD of 0.2to assess the phage load of the seedlings.

## 3. Results

### 3.1 Phage Isolation, Morphology, Annotation and Taxonomy

The novel phages were isolated from winter wheat rhizosphere and wastewater on the campus of the Forschungszentrum Jülich. The *Agrobacterium phage* Alfirin was retrieved from the rhizosphere sample at the IBG-2 crop garden using *Agrobacterium strain C58* as a host. *Pseudomonas phage* Athelas was isolated from a wastewater sampled at Forschungszentrum Jülich wastewater plant using *Pseudomonas syringe pv. lapsa* (DSM 50274)(Figure 1A). Phage Alfirin formed clear plaques with a mean diameter of 0.96 mm. Phage Athelas formed large and clear plaques with an average diameter of 6.86 mm (Figure 1B).

**Figure 1:**
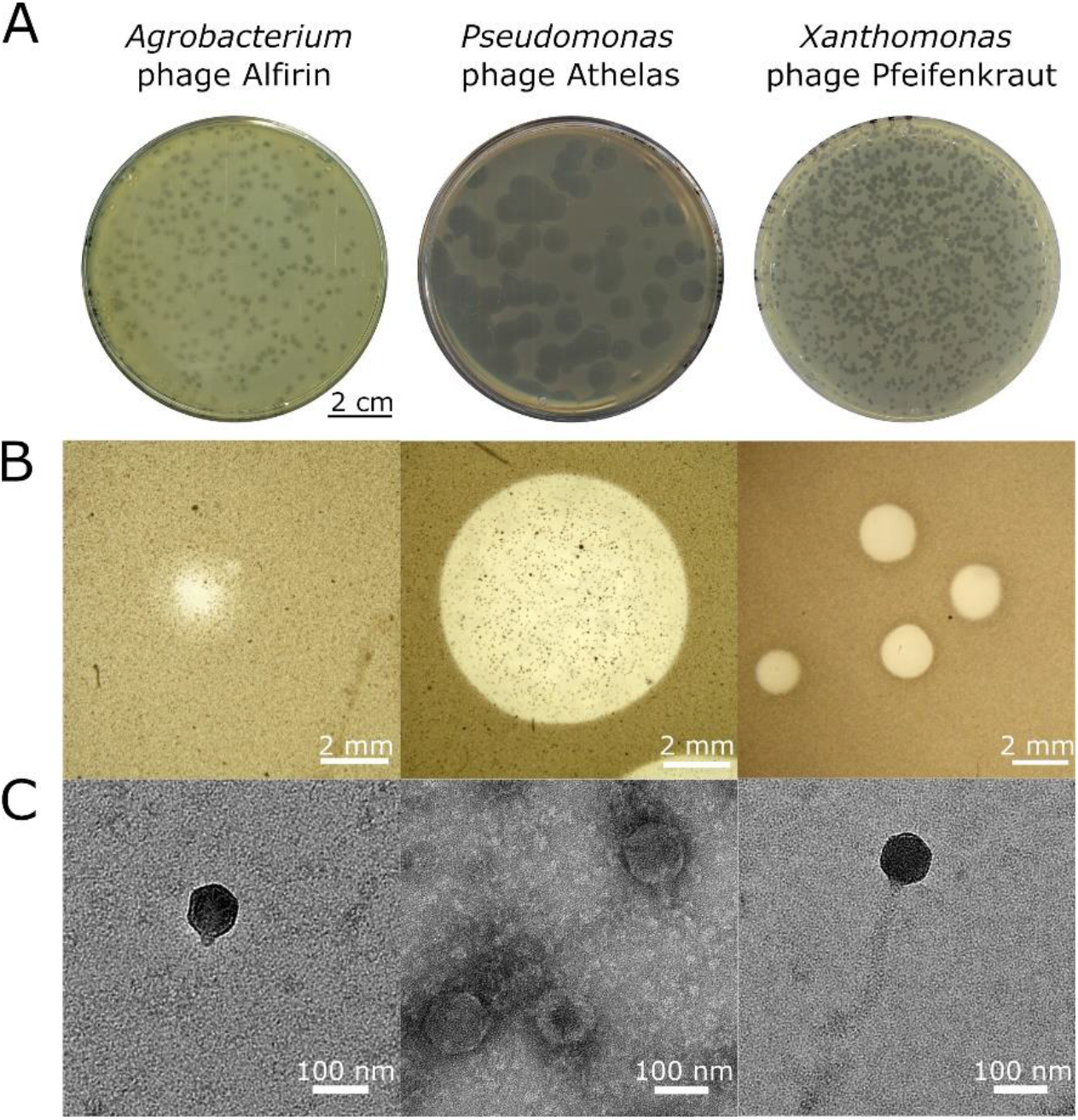
Phage morphology of the novel *Agrobacterium* phage Alfirin, and the *Pseudomonas* phage Athelas as well as *Xanthomonas* phage Pfeifenkraut which was previously described [^32^]. (A) Plaque morphologies of phages on 0.4% soft agar. Scale Bar: 2 cm; (B) Stereo microscopy of single plaques. Scale bar: 1 mm; (C) Transmission electron microscopy (TEM) images of virion particles. The phage isolates were negative stained with uranyl acetate. Scale bar: 100 nm

The isolated phages were sequenced using Illumina MiSeq short-read technology and the genomic features of phage Alfirin, Athelas and Pfeifenkraut are summarized in Table 1, and all other phages used in this study in Table S1. Briefly, the genomes of the novel phages Alfirin and Athelas are 46 and 40 kb in size, with a GC content of 53 and 57%, respectively (Figure S1).

**Table 1.**
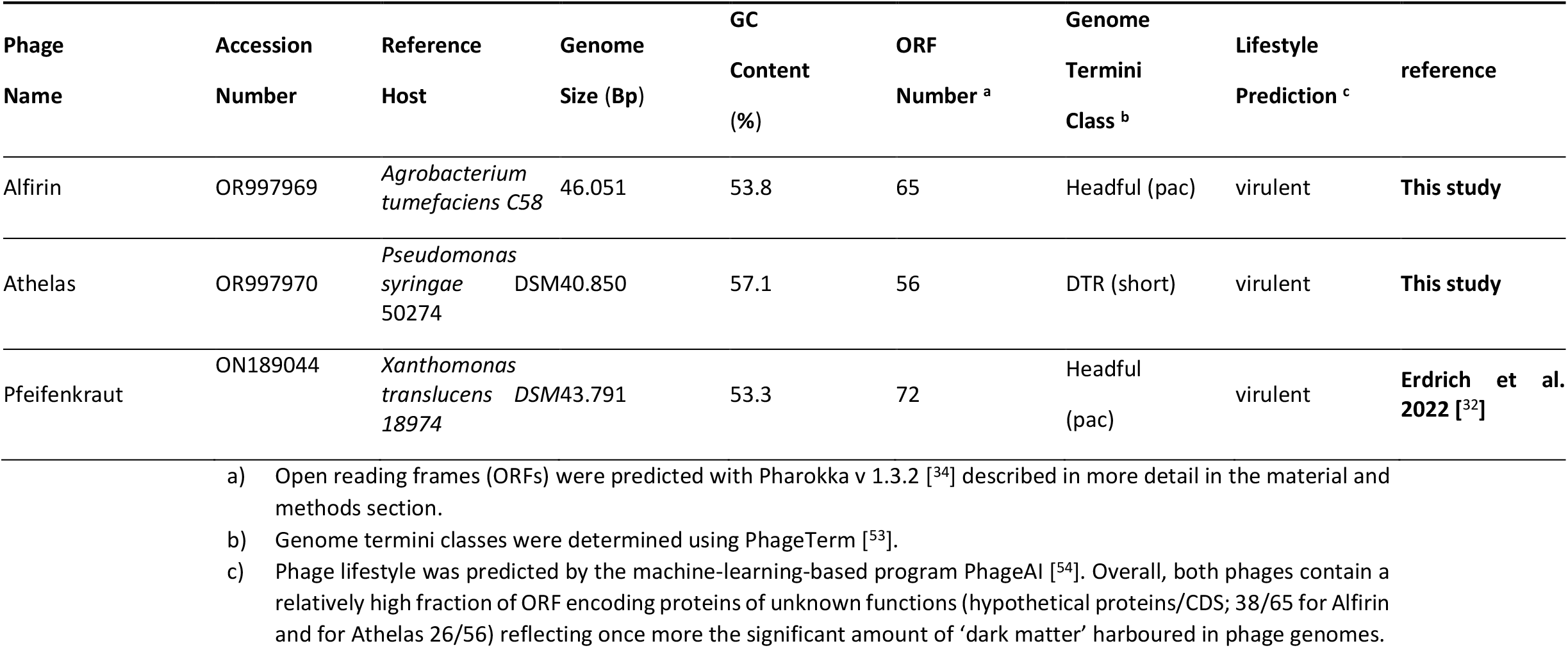
Basic genomic features of Phage Alfirin, phage Athelas and other phages used in this study.

While Alfirin is predicted to follow the headful packaging mechanism[^52^], phage Athelas has short directed terminal repeats (DTRs) of 221 bp. The genomic ends were determined using Phage Term [^53^]. A prerequisite for phage biocontrol is a lytic lifestyle of the bacteriophage, therfore the lifestyle was predicted using PhageAI, a machine-learning tool which compares the genomes to over 20,000 publicly available phages [^54^]. Both newly isolated phages were classified as virulent. This is further supported by the absence of genes coding for an integrase within the genomes.

Comparison of the genomes of our isolates with their closest relatives revealed that phage Athelas is part of a described species and phage Alfirin is its own new species. With an average nucleotide identity of 99% Athelas is a member of the phage NOI species and belongs to the family of Autographiviridae. When compared with the closest relatives phage Athelas clusters with phages isolated on *Pseudomonas syringae pv tomato*. The genome of phage Alfirin shares a 58% sequence identity with *Agrobacterium* phage Atu_02 und therefore forms a new species (Figure S2).

### 3.2 Binding of phages to *Arabidopsis* seeds and influence of the seed coat mucilage

In the following the binding of the newly isolated phages as well as the previously described *Xanthomonas* phage Pfeifenkraut to *Arabidopsis* seeds was investigated.

To test the ability of phages to adhere to seeds we used pre-serilized *Arabidopsis thaliana Col-0* which is a well-known mucilaginous plant [^55^], and has the ability to generate large amounts of seeds in a relatively short time. To discriminate between binding and random co-translocation of the phages on the seeds we washed the seeds twice in ddH_2_O. To detect infectious phage particles bound to *Arabidopsis* seeds, we harnessed one of the hallmarks of phage biology – plaque assays – by placing the seeds after treatment, onto a bacterial lawn containing the respective host species. A visible clearance of the bacterial lawn is consequently indicative for the binding of phage particles to the seed surface [^56^], Figure 2A.

**Figure 2:**
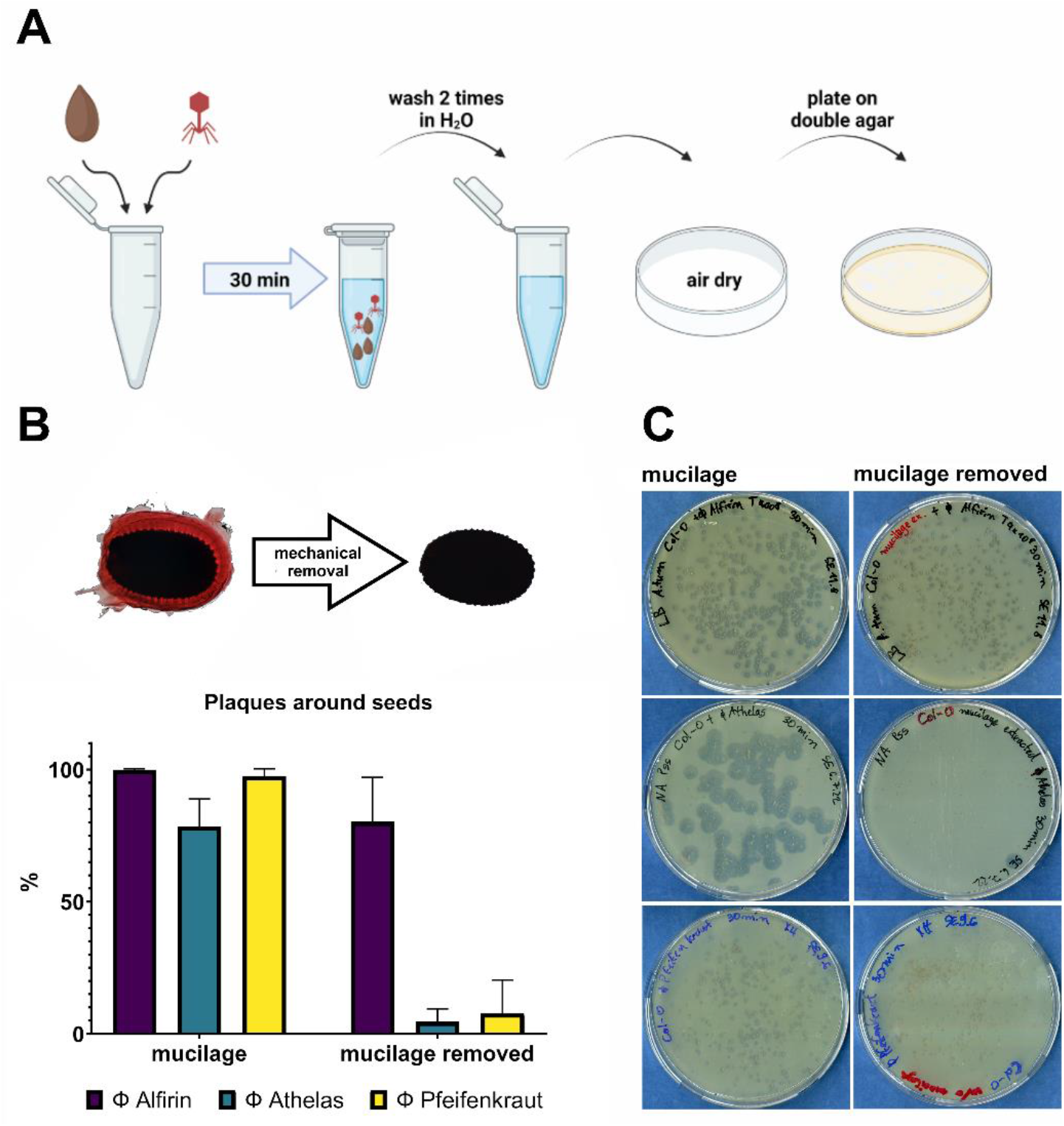
Phage binding to seeds and influence of artificial removal of the seed mucilage. **A)** Seed-coating-workflow; binding of phage particles on seeds of *Arabidopsis thaliana*. **B)** Percentage of plaques detected around seeds for *Agrobacterium* phage Alfirin, *Pseudomonas* phage Athelas or *Xanthomonas* phage Pfeifenkraut incubated on seed with or without mucilage. Wild type Col-0 seed stained with 0.01% ruthenium red solution before and after mechanical removal of the mucilage. **C)** Double agar overlay with phage coated seeds.

After a first observation of phage binding to seeds of *A. thaliana* we asked the question, which mechanism is responsible for binding of the phages. As *Arabidopsis* is described as a mucilaginous plant [^55^], which share the characteristic that the seeds release a matrix of sugars, pectin and cellulose after first contact with water, we tested if the mucilage either structurally or chemically is involved in phage attachment of the seeds. Using non-disturbed seeds and seeds with removed mucilage ([^46^], Figure 2B), we could show that the mucilage is crucial for seed binding for *Pseudomonas* phage Athelas (73% reduction) and for *Xanthomonas* phage Pfeifenkraut (94% reduction) (Figure 2 BC). From this initial set of phages, only phage Alfirin was not significantly depended on the presence of the mucilage.

### 3.3 Phages of the *Autographiviridae* family significantly depend on the presence of the mucilage

In the initial set of phages, phages Pfeifenkraut and Athelas showed a clear dependency on the presence of the mucilage. The strongest effects where reproducibly observed for podovirus Athelas. To test whether this trend holds for morphologically similar viruses, we tested all podoviruses from the BASEL collection [^29^] and also included the model *E*.*coli* phage T7. We could show that T7 as well as Bas64-Bas68, belonging to the *Autographiviridae* family, showed a strong dependence on the mucilage (Figure 3). The podovirus Bas69 belonging to the *Schitoviradae* showed no significant difference in seed adhesion with or without mucilage. These results indicate that the size (surface cross section) alone cannot explain the differences in binding behaviour amongst different phages of similar size. The observed pattern suggests that taxonomically related phages also show similar adhesion properties to the mucilage of plant seeds.

**Figure 3:**
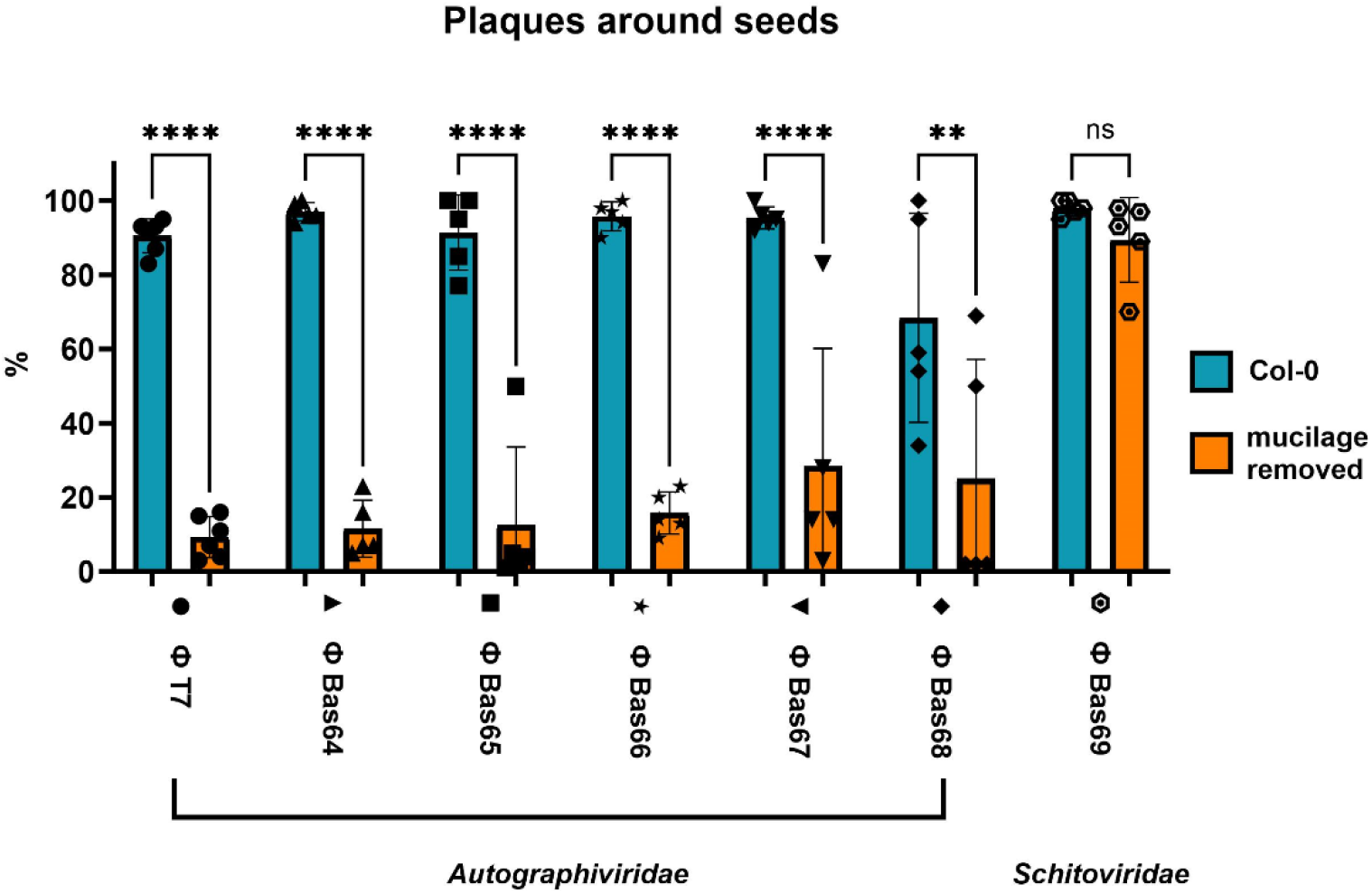
Binding of podovirions to seeds and influence of artificial removal of the seed mucilage. Selected were all BASEL featuring podovirus morphology, as well as model phage T7. Percentage of plaques around seeds with and without mucilage. Each plate contained between 50-300 seeds.

### 3.4 Influence of seed coat mutants on phage binding

To further elucidate which components of the seed coat mucilage are relevant for phage binding we set out to test different *Arabidopsis* seed coat mutants with phage Athelas, because it was affected strongly by presence or absence of the mucilage and produces large plaques enabling robust quantification (Figure 4). As a control, we used phage Alfirin as a control that does not require mucilage. The *rhm2, clsa2-3* and *muci70-1* (Table 2) mutants showed no significant influence on phage binding, indicating that pectin does not seem to be required for phage binding. TRANSPARENT TESTA GLABRA1 (TTG1) is a master regulator involved in many processes and is required for mucilage production. The mutant *ttg1-21* had a significant impact on phage binding for phage Athelas, which confirms the observation that removal of the mucilage impacts phage Athelas binding to seeds. The second strongest impact was observed for the cellulose synthase 5 mutant (*cesa5*) that is required for production of cellulose within the mucilage and for the correct layering of the mucilage [^57^]. Deletion of *cesa5* was reported to cause a reduction of diffusible cellulose within the *Arabidopsis* mucilage. Also, the *subtilisin-like serine proteases 1*.*7* (*sbt1*.*7*) mutant did impact Athelas adhesion to the seed. Hinting in direction of the importance of accessible mucilage sugars for Athelas since sbt1.7-mutant does not release mucilage properly upon hydration [^58^]. Transmission electron microscopy further supported our hypothesis that Athelas directly interacts with polymeric fiber structures within the mucilage of *Arabidopsis* (Figure S3).

**Table 2.** *Arabidopsis* mutants used in this study and their physiological effects.

| Mutant name | Gene name | NASC number | Physiological role | Reference |
| --- | --- | --- | --- | --- |
| *ttg1-21* | transparent testa galbra 1 | GK-580A05 | Master regulator with pleiotropic roles in e.g. trichome initiation, anthocyanin biosynthesis, and seed coat mucilage biosynthesis.<br>Deletion leads to absence of SCM and the name giving transparent testa phenotype. | [59] |
| *sbt1.7-1* | subtilisin-like serine protease 1.7 | n.a | Plays a role in seed mucilage maturation, by degradation of pectin methylesterase inhibitors.<br>Deletion leads to severe mucilage extrusion defect. | [58] |
| *cesa5-1* | cellulose synthase 5 | N2106719 | Cellulase synthase subunit 5 is expressed specific to epidermal cells and coincides with the accumulation of mucilage polysaccharides in the SCM.<br>Deletion leads to repartitioning of mucilage pectin and the absence of diffusible cellulose within the mucilage while the crystalline cellulose is not affected. | [60] |
| *csla2-3* | cellulose synthase-like a2 | SALK_149092 | Is involved in the biosynthesis of mucilage glucomannan and structuring of the crystalline cellulose in the adherent mucilage.<br>Deletion leads to repartitioning of mucilage pectin and the absence of crystalline cellulose rays within the mucilage, while the diffusible cellulose and total sugar content are not affected. | [61] |
| *rhm2/mum4* | rhamnose biosynthesis 2/<br>mucilage modified 4 | SALK_076300 | Required for the synthesis of pectinaceous rhamnogalacturonan I, the mayor component of *A. thaliana* mucilage<br>Deletion leads to reduction of pectin synthesis and thereby to smaller total amounts of mucilage | [62,63] |
| *muci70-1* | Mucilage related 70 | SALK_129524 | Is a pectin related galacturonosyltransferase located in the Golgi apperatus.<br>Deletion leads to reduction of pectin in the SCM, shorter rhamnogalacturonan I (RGI) chains and xylan substitution and over all smaller mucilage capsules around the seeds. | [64] |

**Figure 4:**
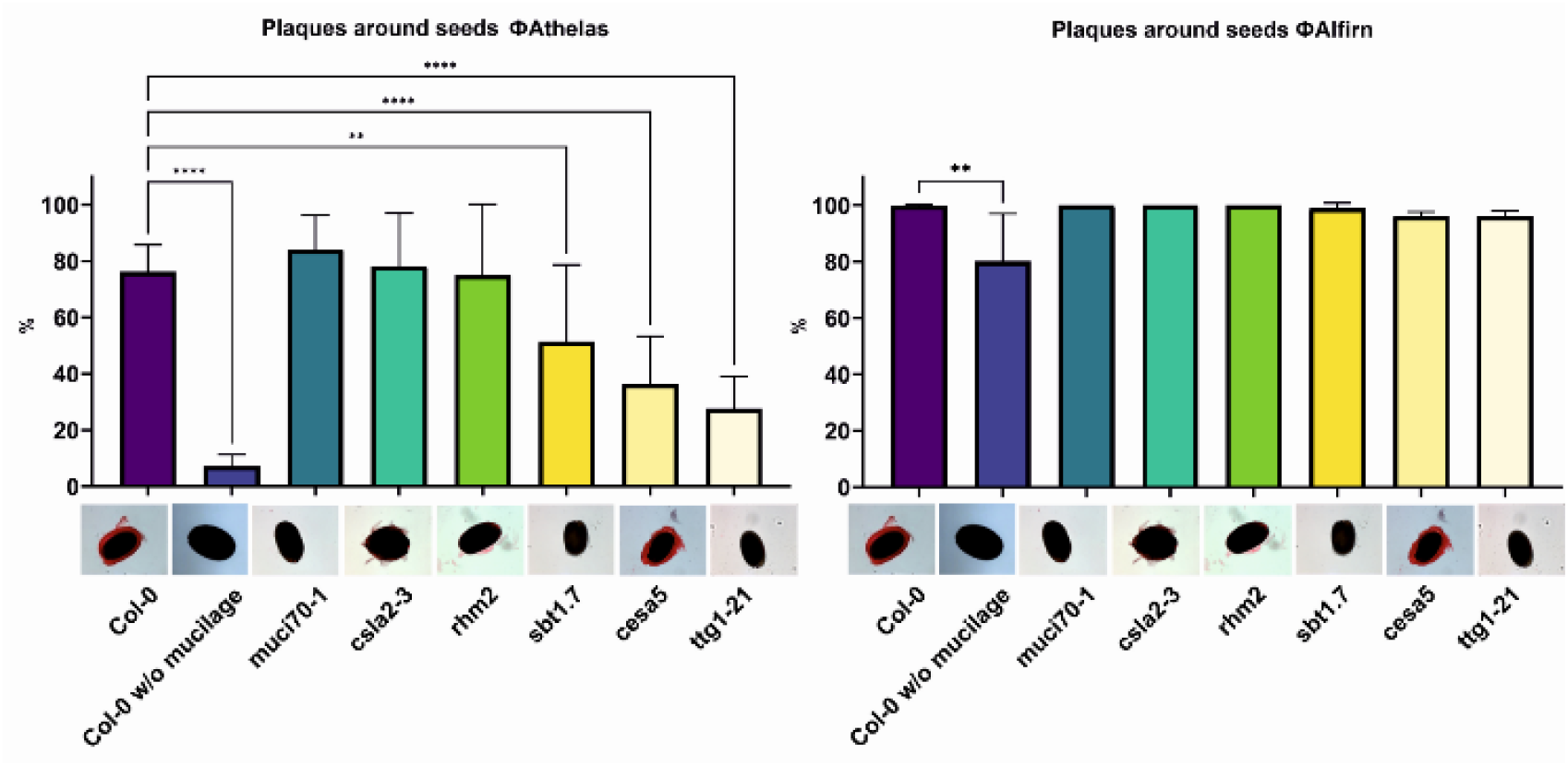
Phage seed coating on different *Arabidopsis* mutants. Left panel phage Athelas; right panel phage Alfirin. Shown is the amount (%) of plaques detected on a double agar overlay post seed coating of the *Arabidopsis* mutants: *muci70-1, csla2-3, rhm2, sbt1*.*7, cesa5* as well as the wild type *A. thaliana (Col-0)* and mechanically removed wild-type seeds. Below each column the respective seed stained with 0.01% ruthenium red solution is depicted.

As expected, phage Alfirin was not affected by the mutants tested in this study. Which is in line with previous experiments showing that the absence of the mucilage did only weakly effect phage Alfirin. Since phages Athelas and phage Alfirin are of similar size the question remains what causes these differences in their binding behaviour to seeds.

### 3.5 Comparative analysis of phage tail fibres

Testing of phage binding to different *Arabidopsis* mutants suggested a relevance of the mucilage polysaccharide fraction, more precisely the diffusible cellulose, on phage binding. From previous studies, it is known that the *E*.*coli Autographiviridae* (T3,T7 and Bas64-Bas68) recognize components of bacterial lipopolysaccharides (LPS) as a receptor [^29,65^]. This hints into direction of a similar mechanism for phage Athelas. To gain more insights into a potential specific, chemical interaction with the seed mucilage, a set of receptor-binding proteins (RBP) from the phages in this study and close relatives was compared phylogenetically and structurally by in silico folding of the proteins (Figure 5). To further investigate the cause of this differential binding behaviour of phages with a similar capsid size as well as a short tail, we compared the host binding proteins of those phages. The phylogenetic tree of the tail fibres revealed that the Basel *Autographiviridae* are a sister group to *Pseudomonas* phage Athelas which was also highly dependent on the mucilage Figure 5 A. The similarity of the tail fibres of this groups can also be seen in the structural 3D model computed with Alphafold (Figur 5B 1 and 2).The *E. coli Autographiviridae* (T3,T7 and Bas64-Bas68) were found to be dependent the bacterial LPS in previous studies [^29,65^]. This hints into direction of a similar mechanism for phage Athelas. Direct sequence comparison revealed that the N-terminal region showed higher conservation between those two regions than the C-terminal fraction. Another interesting observation is that phage Alfirins second tail fibre clusters together with phage Bas69 tail fibre which also was not significantly impacted by the removal of the mucilage.

**Figure 5:**
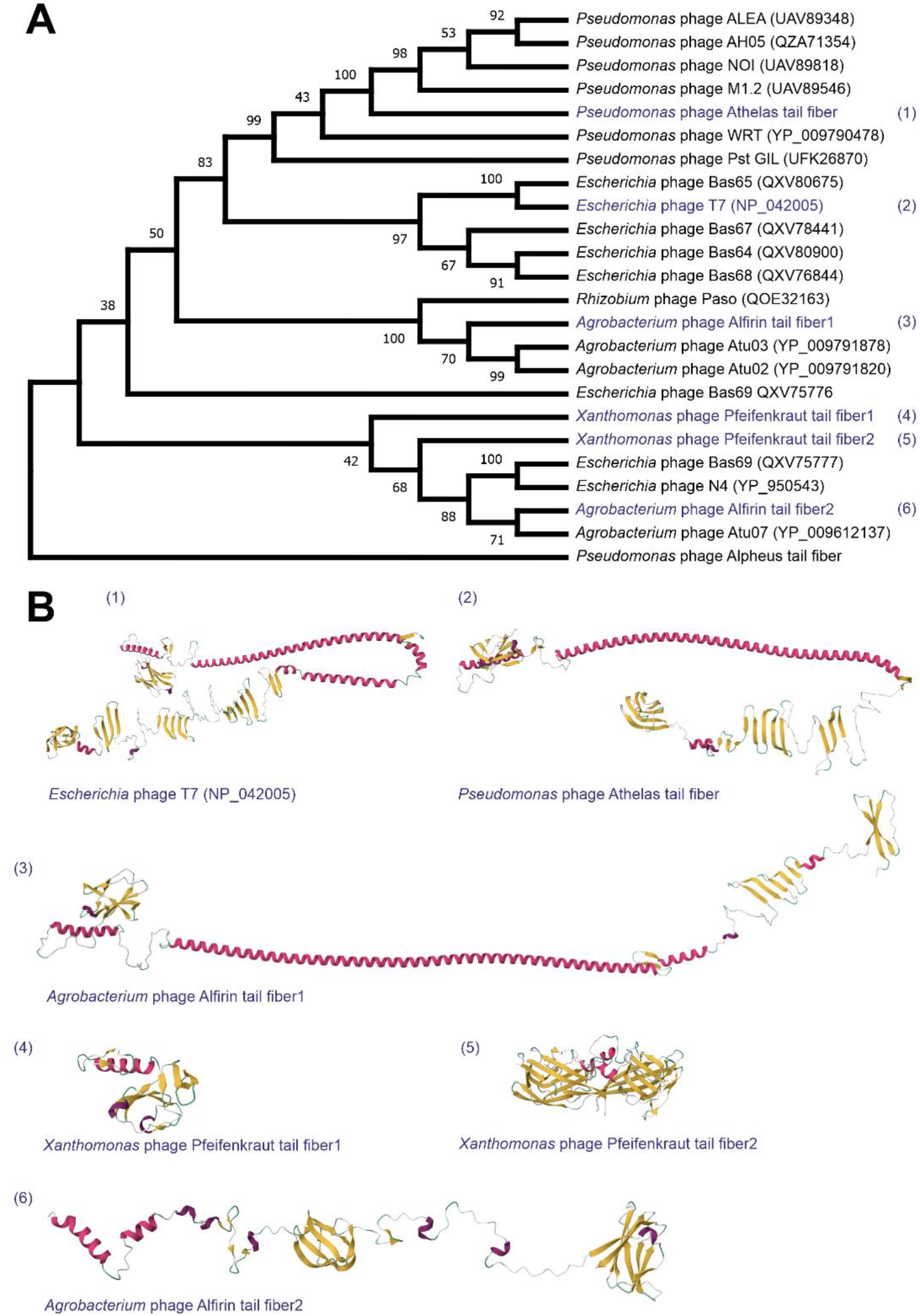
Phylogenetic tree of phage receptor binding proteins and in silico folding. **A)** Phylogenetic tree of phage receptor binding proteins. Ancestral states were inferred using the Maximum Likelihood method [^47^] and JTT matrix-based model [^48^]. Evolutionary analyses were conducted in MEGA11 [^49^]. **B)** Protein structures of selected phage receptor binding proteins. The 3D protein structures were predicted using the ColabFold v1.5.3 webserver [^50^]. N-termini are displayed in the left upper corner of each protein model.

### 3.6 Stability of phages on seed surfaces

A high stability of infectious phage particles on seed surfaces, is prerequisite for the establishment of effective phage-based biocontrol strategies. To test for this, we conducted experiments to determine the storage stability of phages when attached to *A. thaliana* Col-0 seeds. For a time span of more than 4 weeks, phages Alfirin and Pfeifenkraut showed high levels of stability when stored at 4°C. Phage Alfirin showed binding of 99 % and was stable for over 28 days and beyond (Figure 6). A similar pattern was observed for phage Pfeifenkraut with an initial average binding of 96 %. Phage Athelas showed a lower initial binding with 88% and a notable reduction after 14 days.

**Figure 6:**
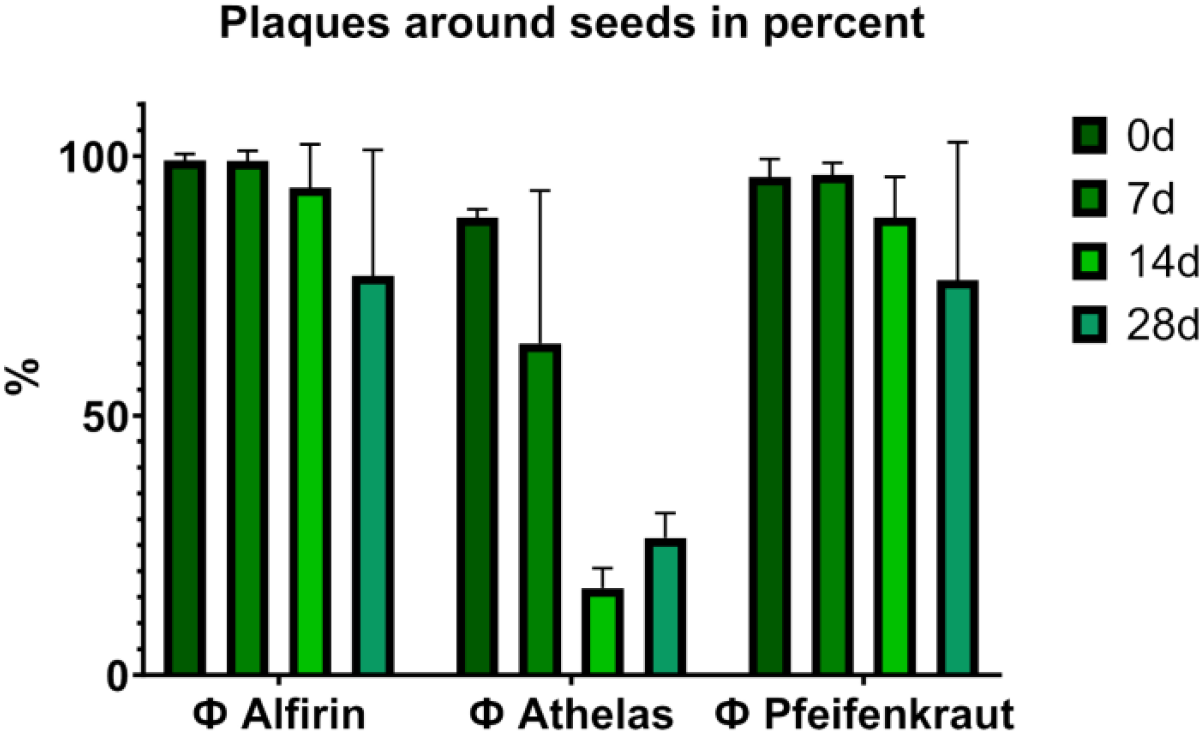
Stability of phage on *Arabidopsis* seeds stored at 4°C. Shown is the percentage of seeds showing plaques when plated on a double agar lawn containing the host bacterium (*A. tumefaciens* for phage Alfirin, *P. syringae* for phage Athelas and *X. translucens* for phage Pfeifenkraut). Portions of 50-200 seed were tested at every time point indicated, for each phage. The experiments were conducted in 3 biological replicates (further controls in phage buffer are shown in Figure S4).

### 3.7 “Boostering” local phage amplification at plant seeds

As a next step, we investigated the possibility to locally increase the amount of infectious phages in planta. By adding a nonvirulent *Agrobacterium* strain without the tumour inducing plasmid (delta Ti) required for infection of the plant [^51^] we aimed at a “boost” of phage production locally without harming the plant. We could show that the survival rate is increased in plants treated with the phage (Booster MOI1) compared to the pathogen condition (Figure 7A). Notably, the plants treated with the phage control (without pathogen) grew even better (Figure 7). The reason from this growth promotion by the phage is so far unclear and will be addressed in future studies. After 14 days in the climate chamber, we checked how many plants still had active phage particles on their surface and therefore transferred the seedlings to a bacterial lawn of *A. tumefaciens* and checked for occurring lysis (Figure 7B). Twice as many plants in the booster condition showed plaques compared to the phage control. This is indicating that the additional delivery of a non-virulent host in combination with the phage can improve phage longevity in planta, and deserves further investigation as an application strategy.

**Figure 7:**
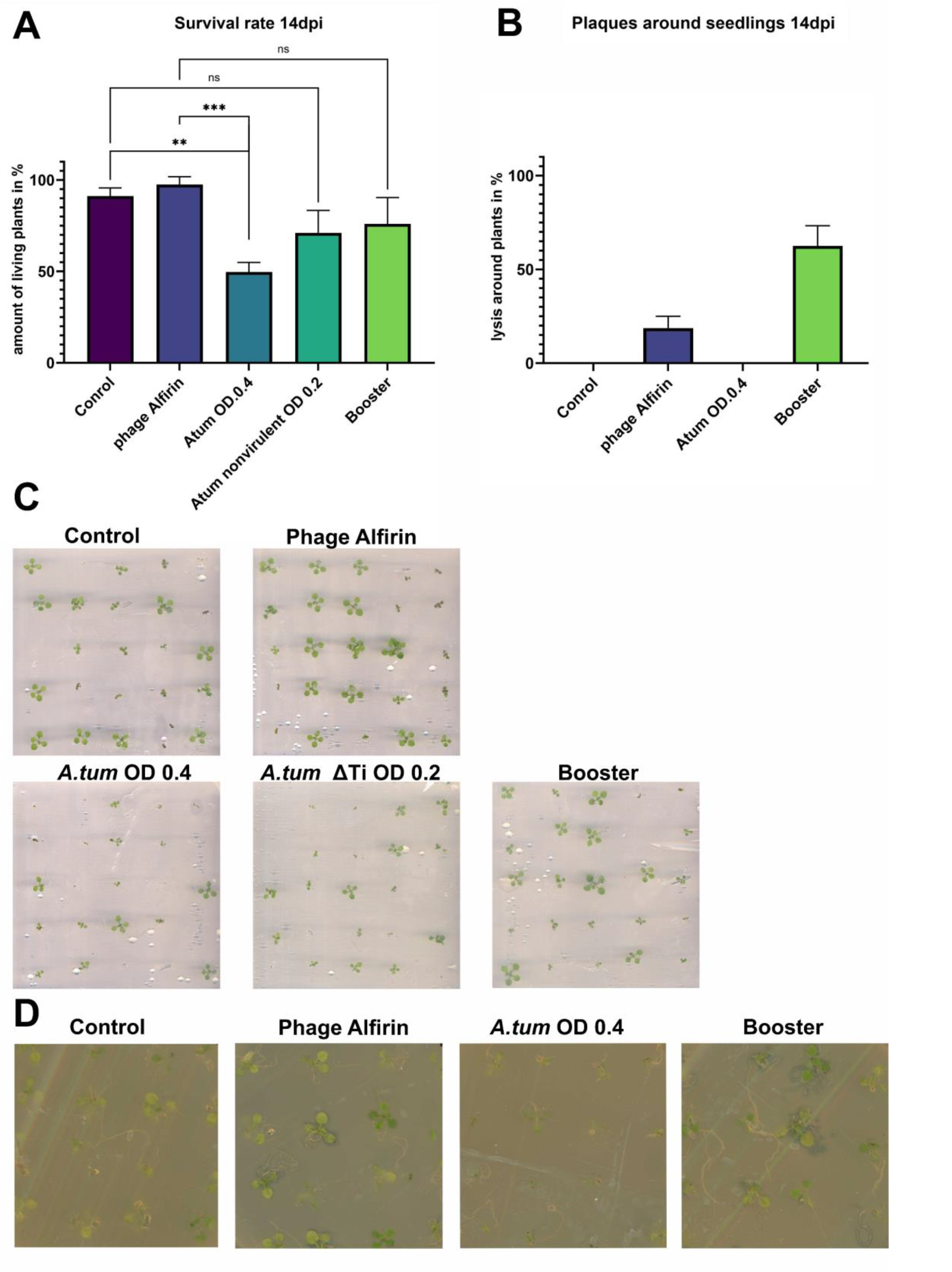
Local phage amplification acceleration with a “booster” that have become integral to the plant-microbiome through co-evolution during the process of plant domestication A Survival rate after 14 days. B Amount of plants with active phages after 14 days. C images of the plants after 14 days. D images of the plants on bacterial lawn

## 4. Discussion

Bacteriophages are a still untapped resource that could advance sustainable biocontrol strategies of plant pathogenic bacteria. This is due to the main characteristics of lytic phages: host specificity and the ability to self-propagate. In this study, we investigated the binding of phages to *Arabidopsis* seeds with a special emphasis on the influence of the seed coat mucilage during this interaction. We confirmed the binding of all phages tested and observed that for some, the seed coat mucilage is crucial for successful seed binding. We linked this dependence to specific mucilage components. Finally, we move towards more application-oriented questions, affirming the stability of phages on mucilage-producing seeds. Additionally, we observed enhanced seed/seedling viability under pathogenic pressure.

The importance of protecting seeds and young plant parts against pathogenic microbes can’t be overstated. In fact, bacterial transmission via seeds was reported with significant yield losses in many cases [^1–8^]. The application of phages as treatment strategy has gained special interest in recent years [^21, 66, 67^]. Successful treatment of plant seeds has been recently demonstrated for *Xanthomonas* (Xcc) in cabbage, for example as a potential treatment in plant nurseries. Here, the authors, showed significant symptom reduction and seed cleaning of artificially contaminated seeds when applying high phage concentrations [^21^]. A further study showed the protection of rice seedlings with phages against *Xanthomonas oryzae* (Xoo), with an emphasis on pre-infection phage treatment, since this showed the strongest protection [^66^]. Apparently, also the pre-treatment of seed tubers increased plant germination, as shown for potatoes [^67^].

Altogether, these studies emphasize the high potential of seed- and pre-treatment strategies, but the mechanisms by which bacteriophages are kept in close proximity to the seeds or young plant parts and their interaction with surface components remain to be understood. In this study we demonstrated that for certain types of phages, the seed coat mucilage has a crucial part in the binding process to the seed. Our experiments have further validated the remarkable stability of infectious phage particles on seed surfaces, extending beyond a period of four weeks, which likely can be further improved by optimizing seed coating formulations. This will open up a variety of options for future applications on seeds that do not produce mucilage naturally.

Mucilage shows multiple independent origins throughout plant evolution [^23^] probably due to it’s functions like maintaining a moist environment for the seedling in a microenviroment, anchorage to soil, increased dispersal [^25^]. On top of that, the mucilage could be also an additional layer of defence against unwanted bacteria, by entrapping bacteriophages in close vicinity of the seeds and root tips. In our study, all tested phages bound to *Arabidopsis* seeds with mucilage. When the mucilage was mechanically removed, phage binding decreased significantly. However, phage specific differences were noted. While phage Athelas and other members of the *Autographiviridae* family showed a very strong dependence on mucilage for binding, phage Alfirin also showed interaction with seed surface in the absence of the mucilage. It would be interesting to investigate if the phage-mucilage dependency emerged as adaptive trait for some phages that have become integral to the plant-microbiome through co-evolution during the process of plant domestication [^68^].

In the study of the adhesion process we differentiated between two mechanisms: (i) adhesion based on a physical structure of the polymer-matrix, where the matrix would function as a mesh with a pore size between 2-50nm [^69^] or (ii) adhesion based on chemical interactions between phages and seed mucilage components. The latter was approached by the systematic testing of seed coat mutants of the model plant *A. thaliana*. Here, phage Athelas showed the strongest effect on the *transparent testing galba 1* (*ttg1*) mutant, followed by the *cellulose synthase 5* (*cesa5*) and *subtilisin protease 1*.7 (*sbt 1*.*7*) mutants. Both the *ttg1* and the *sbt1*.*7* mutants are indicative for the fact that mucilage formation/release is a prerequisite for adhesion of phage Athelas. TTG1 is a master regulator in *Arabidopsis* involved in many processes, including the production of the seed coat mucilage [^70^] and its deletion leads to seeds that produce no mucilage. The deletion of *sbt 1*.*7* is described as a non-release phenotype, because it is needed for the regulation of pectin methylesterases crucial for mucilage release in *A. thaliana* seeds[^58^]. Most interestingly *cesa5* which is still releasing the mucilage, but possesses less diffusible cellulose in the mucilage [^60^], shows a significant reduction in binding of phage Athelas. This result suggested that phage Athelas requires this diffusible cellulose fraction for binding to the seed surface. This interaction could potentially be based on the attachment of phage tail fibres to the glucose units of the diffusible cellulose. Further evidence for this hypothesis is supported by the fact many bacterial genes are capable to produce cellulose as part of their biofilm and phages interact with them [^71^]. This was also reported for the plant pathogen *Pseudomonas syringae* [^72,73^]. Nevertheless, further studies will have to test whether this hypothesis holds true and to identify the specificity determinants for this transient interaction.

While phages Alfirin and Pfreifenkraut showed high stability on seeds, it remains unclear why the infectivity of phage Athelas dropped drastically on seeds already after 14 days of storage, while it showed high stability in phage buffer (Figure 3). One possible explanation could be that the mucilage-polysaccharides are able to trigger the DNA ejection. A similar observation was described for bacterial LPS triggered release of the phage genome in Podovirions [^74–76^]. Another possibility could be that the drying process impacts phage Athelas stability, which could be theoretical overcome by adding stabilizers used in classical phyllosphere-phage formulations [^77^]

Stability and titer of phages on seed surfaces can certainly be improved by optimizing seed coating formulations. Here, knowledge gained regarding the specificity determinants of chemical interactions will provide a powerful basis to improve the composition of seed coatings. This could be especially useful for plants that do not produce mucilage naturally. First attempts in to the direction of artificial seed coating with chemical formulations have been reported, e.g. in maize chemical deployment of phages with polyvenylalcohol [^78^]. Further, the application of non-virulent host species could serve as a way to amplify the effective phage titer in the proximity to the plant and thereby enhance protection.

On the larger scale, while we are confident that phages show potential as biocontrol agents and have clarified part of the mechanism for successful phage application to seeds, we also observed that phage-coated seeds germinated without pathogen pressure show a trend to improved germination. This opens the question whether the use of phages as plant protection would also positively influence plant growth and which are the underlying mechanisms.

In summary, the results reported in this study show effective binding of phage binding to *Arabidopsis* seeds and further emphasized that some phages, particularly podoviruses belonging to the *Autographiviridae* are strongly dependent on chemical interactions with the seed coat mucilage. Phage based biocontrol on the seed level certainly has a great potential for application. The chemical universality of some carbohydrates present in bacterial lipopolysaccarides might allow the targeted binding of phages to plant surfaces displaying similar sugar moieties. The better understanding of the molecular basis for these transient interactions therefore has a high potential of the establishment of targeted phage delivery strategies with a high relevance for applications in agriculture and medicine. Further clearing of seeds with phages was shown to be an effective strategy for selective seed cleaning against pathogenic bacteria, leaving the beneficial microbiota intact.

The ecological significance of our discovery that mucilage can bind phages raises important questions that merit further investigation. Is this binding merely coincidental, or is there a conserved chemical nature in the mucilage-microbe interface across different kingdoms of life? Would this also lead to a transmission of phages from plants to the next generation? These questions are certainly highly relevant in the context of plant-microbe interactions but also in the context of effective phage-based biocontrol strategies.

## Author Contributions

Conceptualization, All; methodology, All; validation, All; data analysis, All; investigation, S.E.; resources, B.A, U.S and J.F.; data curation, S.E and writing—original draft preparation, S.E.; writing—review and editing, All; visualization, S.E. supervision, U.S., J.F. and B.A.; All authors have read and agreed to the published version of the manuscript.

## Acknowledgments

The authors gratefully acknowledge the electron microscopy training and access time granted by the biological EM facility of the Ernst-Ruska Centre at Forschungszentrum Jülich. Further, the authors thank Prof. Dr. Franz Narberhaus (RUB, Bochum) for providing both *Agrobacterum tumefaciens* C58 strains.

