## Supplementary material for "Seed coating with phages for sustainable plant biocontrol of plant pathogens and influence of the seed coat mucilage": Figure S1-4 and Table S1

A

### *Agrobacterium phage Alfirin*

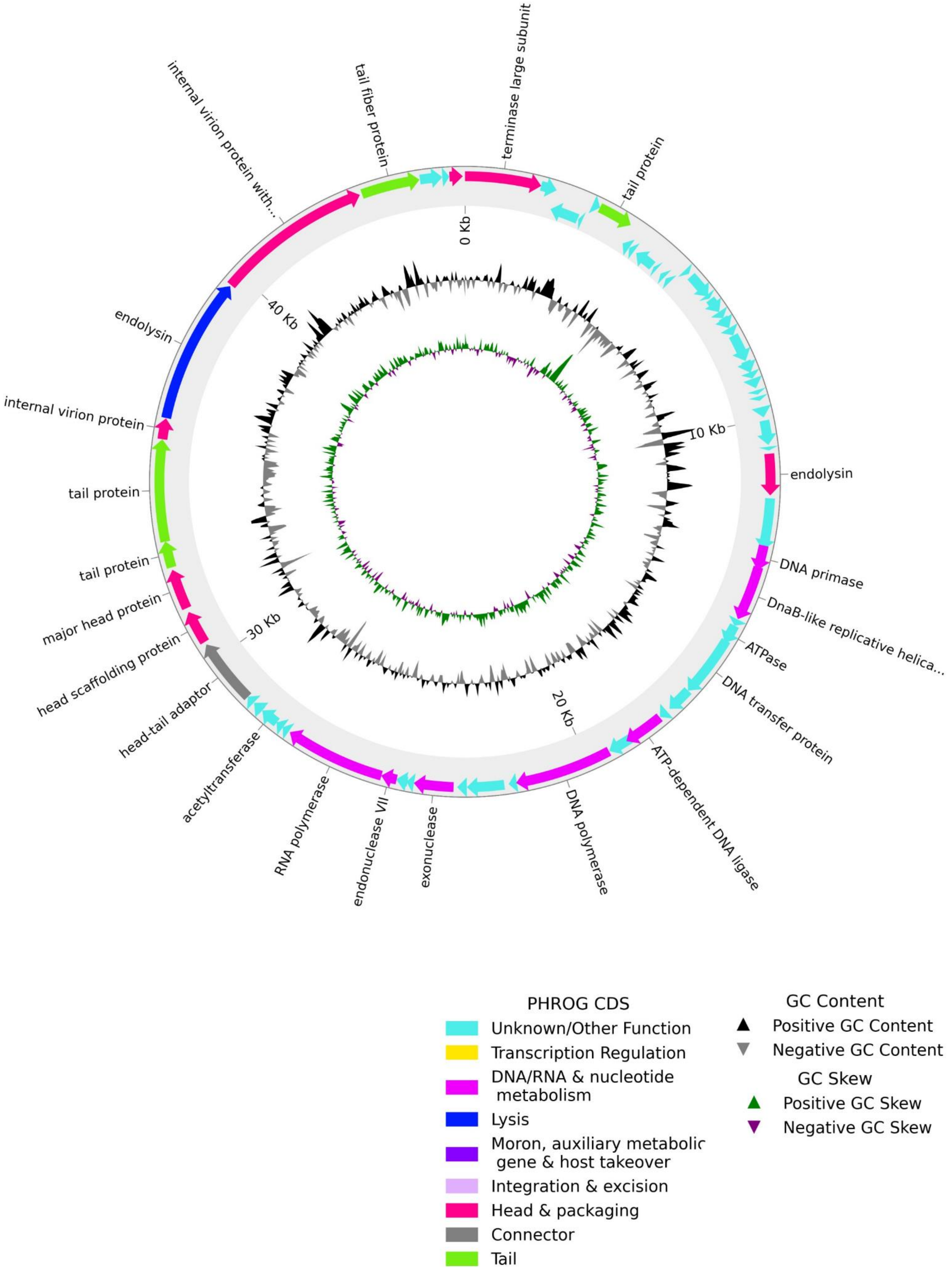

**B** ***Pseudomonas phage Athelas***

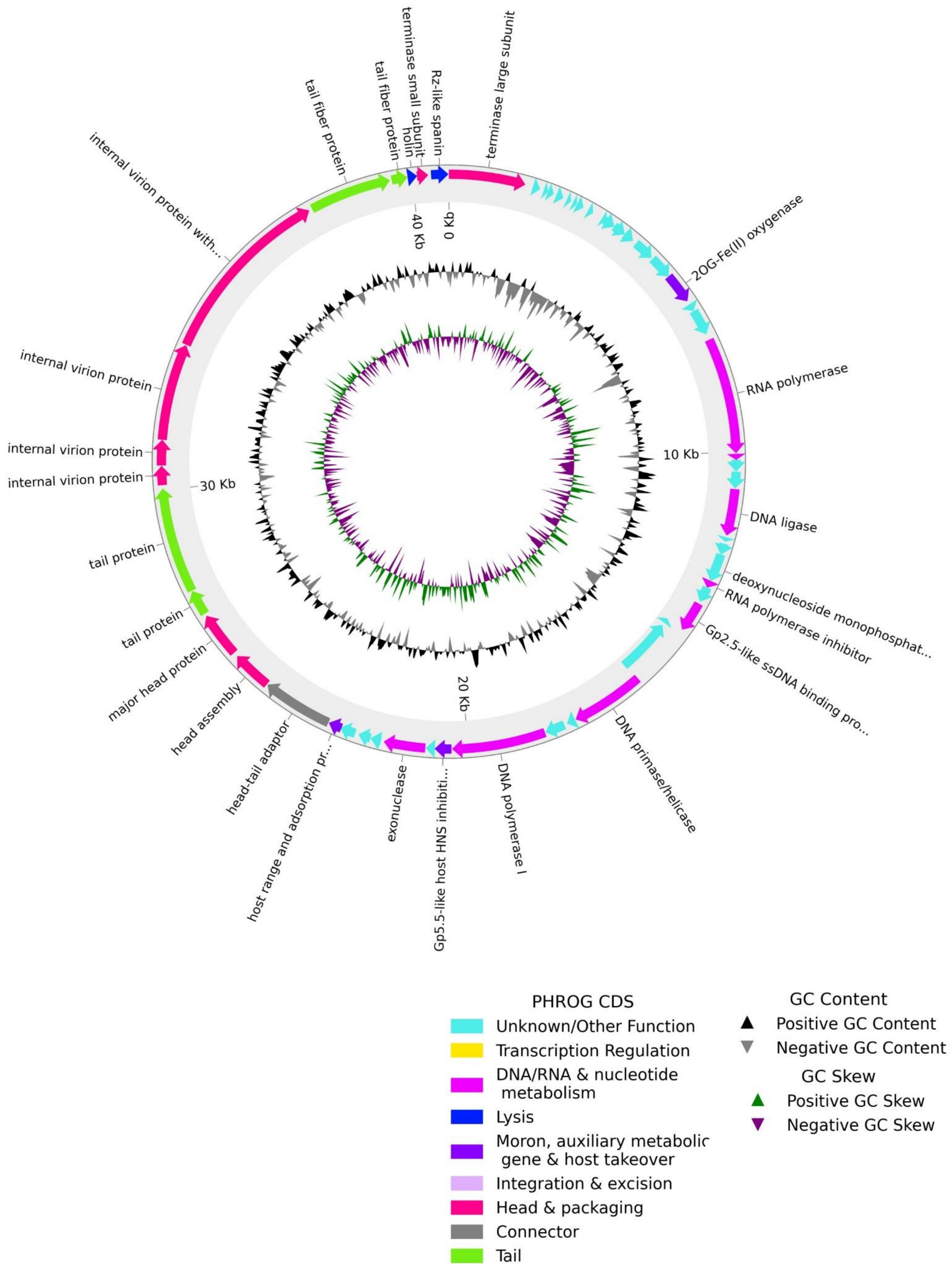

**Figure S1: Phage Annotation of *Agrobacterium* Alfirin and *Pseudomonas* phage Athelas.** A) genome annotation of *Agrobacterium* Alfirin B) genome annotation of *Pseudomonas* phage Athelas The phage open reading frames (ORFs) were predicted with Pharokka v 1.3.2 [34] in terminase reorientation mode using PHANOTATE [35]. Functional annotation was generated by matching each CDS to the PHROGs [39], VFDB [40] and CARD [41] databases using MMseqs2 [42] and PyHMMER [43]. Contigs were matched to their closest hit in the INPHARED database [44] using mash [45]. Plots were created with the pyCirdizen package. The architecture of the genomes of the phages is notably characteristic, as functional units tend to cluster together, demonstrating the inherent modularity of phage genomes.

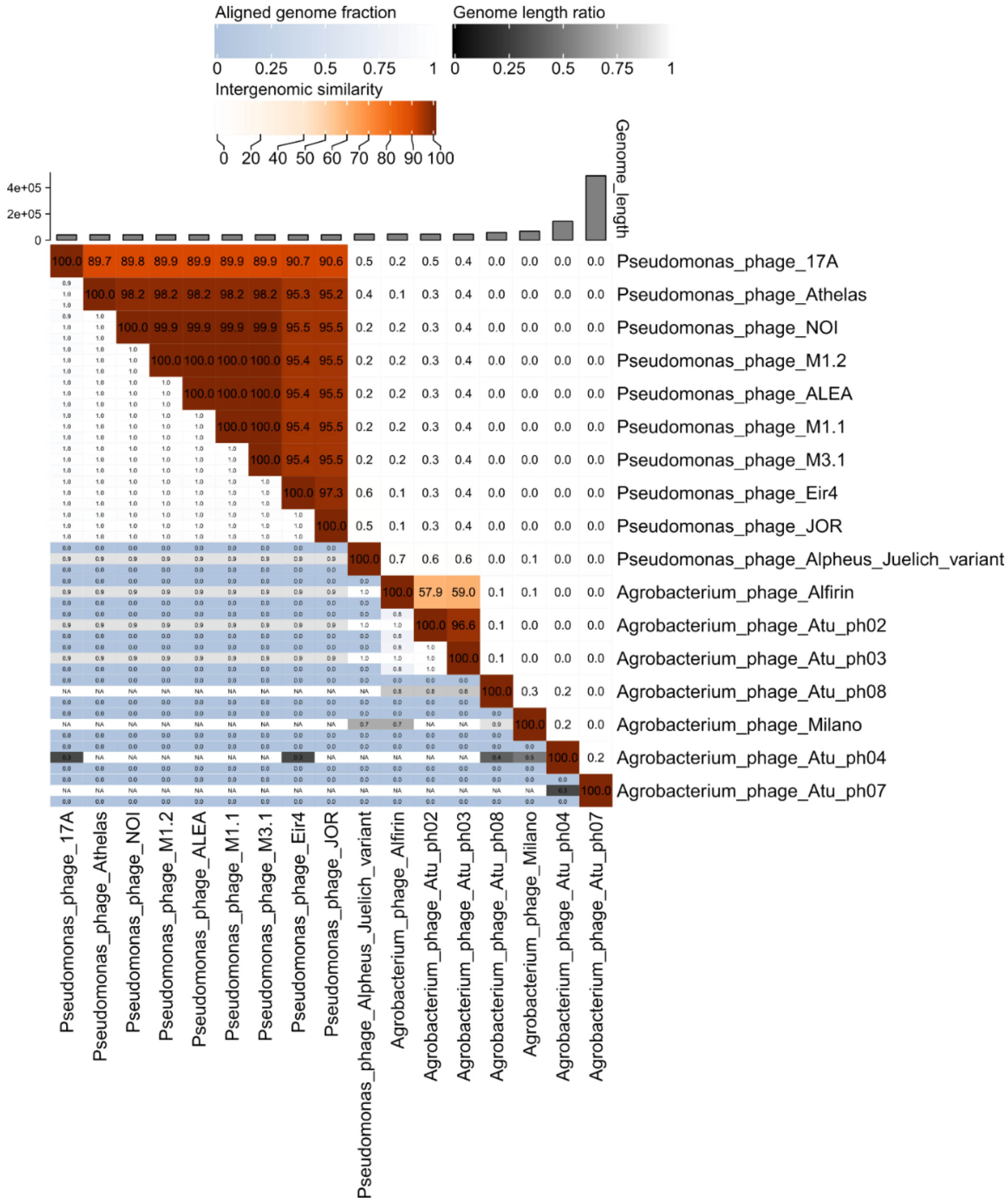


**Figure S2: Viridict Heatmap.** Average nucleotide-based heatmap analysis using the *Agrobacterium* Phage Alfirin and *Pseudomonas* phage Athelas. Genomes are were acquired from NCBI based on relatedness to the isolated phages in this study phages. The analysis was performed using VIRIDICT [80].

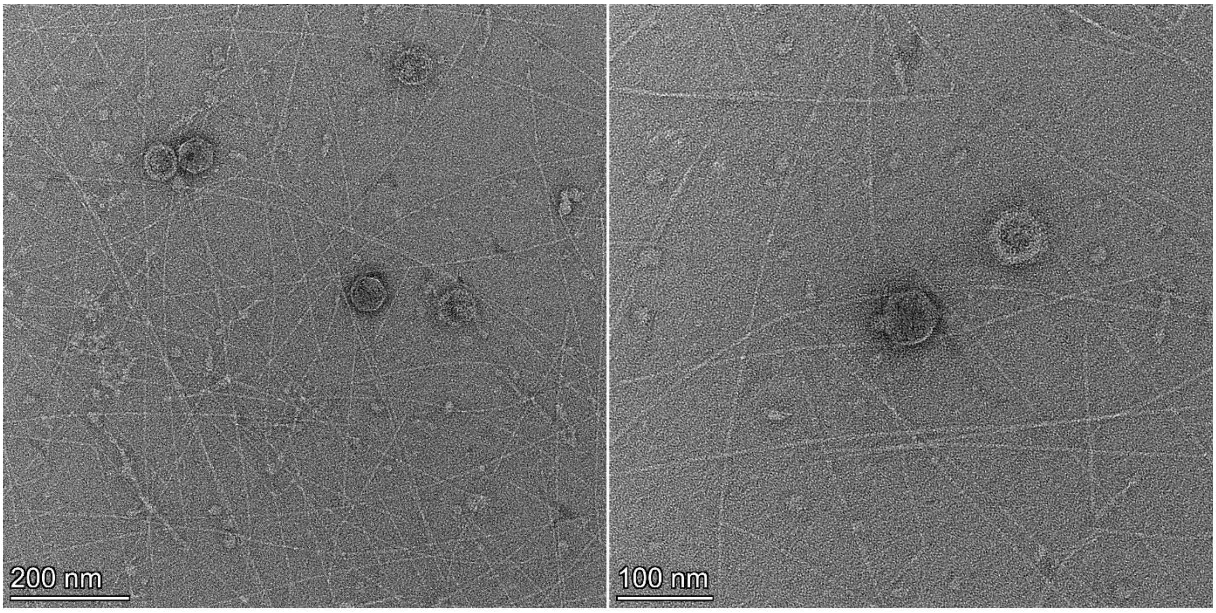


**Figure S3: Electron microscopy images of phage Athelas co-incubated with *A. thaliana* mucilage.** Transmission electron microscopy (TEM) images of phage Athelas virion particles, incubated with mucilage mechanically removed from 2000 Col-0 seeds. After an incubation time of 1h the phage particles were negative stained with uranyl acetate.

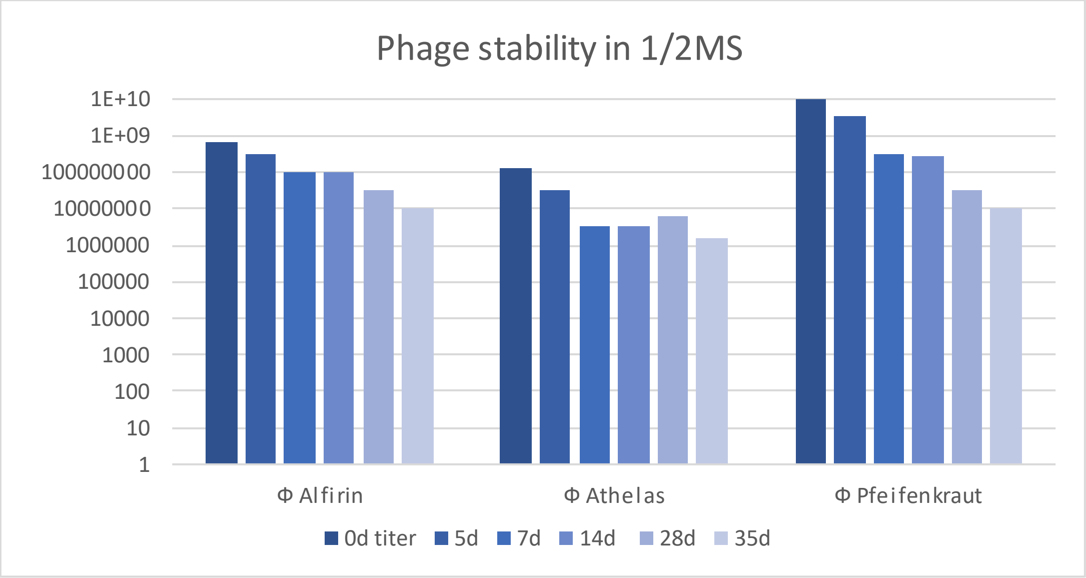


**Figure S4. Phage stability in SM buffer at 4°C.**

**Table S1: Genetic features of the used *E.coli* phages.**

| **Phage Name** | **Accession Number** | **Reference Host** | **Genome Size (Bp)** | **GC Content (%)** | **ORF Number [a]** | **Genome Termini Class [b]** | **Lifestyle Prediction [c]** | **reference** |
|---|---|---|---|---|---|---|---|---|
| Bas64 | MZ501081 | *Escherichia coli K-12* | 39.842 | 48.0 | 51 | n.a. | virulent | **Maffei et al 2021 [[29]]** |
| Bas65 | MZ501078 | *Escherichia coli K-12* | 39.451 | 49.0 | 50 | n.a. | virulent | **Maffei et al 2021 [[29]]** |
| Bas66 | n.a. | *Escherichia coli K-12* | n.a. | n.a. | n.a. | n.a. | virulent | **Maffei et al 2021 [[29]]** |
| Bas67 | MZ501064 | *Escherichia coli K-12* | 39.315 | 49.0 | 48 | n.a. | virulent | **Maffei et al 2021 [[29]]** |
| Bas68 | MZ501055 | *Escherichia coli K-12* | 39.466 | 49.0 | 49 | n.a. | virulent | **Maffei et al 2021 [[29]]** |
| Bas69 | MZ501049 | *Escherichia coli K-12* | 70.849 | 41.0 | 86 | n.a. | virulent | **Maffei et al 2021 [[29]]** |
| T7 | NC_001604 | *Escherichia coli* | 39,739 | | 59 | DTR | virulent | **Studier 1972 [[81]]** |
